# Splicing impact of deep exonic missense variants in *CAPN3* explored systematically by minigene functional assay

**DOI:** 10.1101/2020.03.26.009332

**Authors:** Eugénie Dionnet, Aurélia Defour, Nathalie Da Silva, Alexandra Salvi, Nicolas Lévy, Martin Krahn, Marc Bartoli, Francesca Puppo, Svetlana Gorokhova

## Abstract

Improving the accuracy of variant interpretation during diagnostic sequencing is a major goal for genomic medicine. In order to explore an often overlooked splicing effect of missense variants, we developed the functional assay (“minigene”) for the majority of exons of *CAPN3*, the gene responsible for Limb Girdle Muscular Dystrophy (LGMD). By systematically screening 21 missense variants distributed along the gene, we found that eight clinically relevant missense variants located at a certain distance from the exon/intron borders (deep exonic missense variants) disrupted normal splicing of *CAPN3* exons. Several recent machine learning based computational tools failed to predict splicing impact for the majority of these deep exonic missense variants, highlighting the importance of including variants of this type in the training sets during the future algorithm development. Overall, 24 variants in *CAPN3* gene were explored, leading to the change in the ACMG classification of seven of them when results of the “minigene” functional assay were taken into account. Our findings reveal previously unknown splicing impact of several clinically important variants in *CAPN3* and draw attention to the existence of deep exonic variants with a disruptive effect on gene splicing that could be overlooked by the current approaches in clinical genetics.

## INTRODUCTION

Limb Girdle Muscular Dystrophy 2A or R1 (LGMD2A, LGMDR1; OMIM# 253600), the most frequent form of Limb Girdle Muscular Dystrophy (LGMD) worldwide, is an autosomal recessive disorder characterized by muscle weakness affecting predominantly proximal limb muscles, elevated serum creatine kinase and necrosis/degeneration process observed in muscle biopsy (Fardeau et al. 1996). This disease is caused by pathogenic variants in *CAPN3* (OMIM# 114240, 15q15.1, NM_000070.2), a gene coding for 94 kDa protein calpain3 (Calcium-dependant papain-like protease, P20807) that is a muscle-specific member of the calpain family of calcium-dependent enzymes. Calpain3 acts in muscle sarcomere formation and remodeling (Duguez, Bartoli, and Richard 2006). *CAPN3* is the most commonly mutated gene in patients presenting with Limb Girdle Muscular Dystrophy (Duno et al. 2008), making it one the first genes to be sequenced as part of the diagnostic work up of patients with this type of myopathy (Krahn et al. 2019). More than 400 unique pathogenic or likely pathogenic variants have been reported in *CAPN3* gene so far (359 in LOVD (accessed December 2, 2019) and 555 variants in ClinVar (version November 27, 2019)). However, given the large size of the gene (24 exons, 3316nt), previously unknown variants in *CAPN3* are still being identified in diagnostic laboratories. Protein truncating variants (PTV) in *CAPN3*, such as frame-shift inducing indels, nonsense variants or variants disrupting canonical splicing sites (+/−2 nucleotides from exon-intron junctions), are generally accepted as pathogenic loss-of-function, especially if calpain3 protein was absent in Western Blot performed on muscle biopsy sample. Assigning the clinical significance to a newly identified exonic single nucleotide variant (SNV, either synonymous or missense) is much more difficult, despite help from ACMG guidelines that take into account several lines of pathogenicity evidence (Richards et al. 2015). It is important to note that synonymous variants or missense variants with low impact on protein function can still be pathogenic by disrupting splicing (Duguez, Bartoli, and Richard 2006; Kergourlay et al. 2014). There is a great risk of misinterpreting the clinical significance of these variants using standard approaches, which could potentially lead to incorrect diagnosis. Unfortunately, it is not currently known if certain exonic regions are more likely to harbor splicing-affecting variants. Even though a number of algorithms predicting the splice effects of variants are now available (Rowlands, Baralle, and Ellingford 2019), often it is difficult to estimate their validity in the absence of functional analysis.

In order to identify exonic single nucleotide variants with effect on splicing and to improve the diagnostics of LGMDR1, we developed systematic functional cell-based assay (Minigene) for the majority of exons in *CAPN3* gene. We then selected a representative set of missense variants located outside of exon/intron junction sites (deep exonic variants) and tested the effect of these variants on splicing using the developed Minigene assay. We observed that eight out of 21 selected deep exonic missense variants induced abnormal splicing, leading to a change in the classification of clinical significance for half of them. Interestingly, the majority of the deep exonic variants impacting splicing were not identified by several splice-prediction algorithms tested, highlighting the critical need for robust methods of functional analysis of putative variants disrupting splicing. Thus, in addition to the direct benefit for diagnostics of LGMD2R1 patients, our study draws attention to the existence of deep exonic variants with a disruptive effect on gene splicing that could be overlooked by the current approaches in clinical genetics.

## MATERIALS AND METHODS

### Functional splicing assay for *CAPN3* gene

Minigene reporter assay (Gaildrat et al. 2010) was developed for 18 *CAPN3* exons. Since this assay is not adapted for U12 type introns, The Intron Annotation and Orthology Database (IAOD)(Gault et al. 2017; Turunen et al. 2013) was used to identify this type of introns, leading to exclusion of exons 19 and 20 from the analysis. As described previously (Kergourlay et al. 2014; Puppo et al. 2015), exons and approximately 150 bp of flanking introns were amplified using the Expand high fidelity PCR system (Roche, Basel, Switzerland). Amplicons were subsequently cloned into the pCAS2 vector. PCR and digestion product purifications were performed using the NucleoSpin Gel and PCR clean Up (Macherey Nagel, Dürel, Germany), ligations by Quick ligation kit (New England Biolabs, Ipswich, MA, USA), and transformations in 10-beta Electrocompetent E. coli (New England Biolabs, Ipswich, MA, USA). Variants were introduced into the wild type constructions using the Quick-change II XL site-directed mutagenesis kit (Agilent, Santa Clara, CA, USA). Presence of an abnormally spliced transcript associated with the decrease of the normal transcript was considered as “Impact on splicing”. If an abnormally spliced transcript was present but no decrease of the normal transcript was observed for the mutated construct compared to control, the minigene assay conclusion was “Mild impact on splicing”.

### Cell culture and transfection

C2C12 cells were cultured in DMEM, high glucose, GlutaMAX supplemented with pyruvate, 10% Fetal bovine serum (Life technologies, Carlsbad, CA, USA) and 1% antibiotic and antimycotic. Transfections were made with Fugene HD (Promega, Madison, WI, USA) according to the manufacturer’s instructions.

### Transcriptional study

RNAs were isolated 48 hours after cell transfection, using Trizol/chloroform and DNA Free Removal kit (Invitrogen, Carlsbad, CA, USA). RNAs were reverse-transcribed into cDNA and amplified using SuperScript™ III One-Step RT-PCR System with Platinum™ Taq DNA Polymerase (Invitrogen, Carlsbad, CA, USA). PCR amplifications were performed using specific primers located in exon A and exon B of pCAS2. PCR products were separated by electrophoresis in a 2% agarose gel stained with 0.5 μg/ml of ethidium bromide. A purification step from the agarose gel was performed when several transcripts were present in order to subclone each of them into pGEM®-T Easy Vector System I (Promega, Madison, WI, USA) and analyze them separately. Finally, sequencing of each transcript was performed using the Big DyeR Terminator V1-1 Cycle Sequencing Kit (Life technologies, Carlsbad, CA, USA) on ABI Prism 3130xl capillary DNA Sequencer.

### Variant selection, annotation and classification

*CAPN3* transcripts ENST00000397163 and NM_000070.2 were used for all analyses. 328 unique missense *CAPN3* variants were downloaded from LOVD website for *CAPN3* gene (https://databases.lovd.nl/shared/genes/CAPN3, accessed on December 2, 2019). 286 unique missense variants in *CAPN3* present in ClinVar database were downloaded using Simple CinVar website (http://simple-clinvar.broadinstitute.org/, version of ClinVar database: November 27, 2019, Perez-Palma2019). 447 *CAPN3* missense variants found in the general population were downloaded from gnomADv.2.1.1 (Karczewski et al. 2019) using UCSC table browser (Karolchik et al. 2004). Only missense variants (i.e. exonic non-synonymous non-protein truncating single nucleotide substitutions) were included in the analysis. *CAPN3* missense variants from LOVD were assigned the “Clinical Significance” values based on the classification system used in ClinVar (uncertain/conflicting, likely pathogenic, pathogenic/likely pathogenic or pathogenic). We combined missense *CAPN3* variants from LOVD and ClinVar annotated as non-benign in both databases, producing a set of 403 variants. Of these, 381 variants were located more than 1 or 2 nucleotides away from exon-intron junction. The clinical significance for LOVD variants were then expressed in the same terms as for ClinVar variants. The summary of the variants from this set are shown in the Supplemental Figure 1. Two novel *CAPN3* variants from LGMDR1 cohort at the Department of Medical Genetics at the Timone Hospital (Marseille, France), as well as four previously reported variants, were deposited in the LOVD database (Fokkema et al. 2011) as described in Supplementary Table 1. A representative set of 21 missense *CAPN3* variants located across 16 exons was selected for functional analysis by minigene (Figure 1, Table 1). These variants were located outside of the canonical splice sites (+/− 2nt from exon junctions), were not previously known to disrupt splicing, but were predicted to have an effect on splicing according to Human Splice Finder (HSF version 3.0, http://www.umd.be/HSF3 (Desmet et al. 2009). Three additional variants with uncertain pathogenicity classification were also tested by minigene approach: two variants located at the canonical splice junction sites c.498G>A, p.(Gln166=) and c.1913A>C, p.(Gln638Pro) as well as one synonymous variant c.984C>T, p.(Cys328=) (Supplementary table 1). The REVEL (Ioannidis et al 2016) and CADD scores (Rentzsch et al. 2019) for all the variants in this study were obtained using Variant Ranker (http://vsranker.broadinstitute.org/, (Alexander et al. 2017)). Variant impact of calpain3 protein structure was assessed using VarMap web tool (Stephenson et al. 2019). SpliceAI scores for the variants were obtained using SpliceAI tool (v1.3, https://github.com/Illumina/SpliceAI (Jaganathan et al. 2019)). Effect on splicing predictions were also done using MMSplice (Cheng et al. 2019). SCAP scores were downloaded from http://bejerano.stanford.edu/scap/ (Jagadeesh et al. 2019). MutPred Splice annotation was done using http://www.mutdb.org/mutpredsplice (Mort et al. 2014). MaxEntScan scores (Yeo and Burge 2004) were obtained using http://www.umd.be/HSF3 (Desmet et al. 2009). Variants with MaxEnt threshold score of 3 or a score difference of more than 30% with the wild-type score were considered as splice disrupting. For 13 out of 24 variants tested, the output was “No result found with this matrix”.

**Figure 1:**
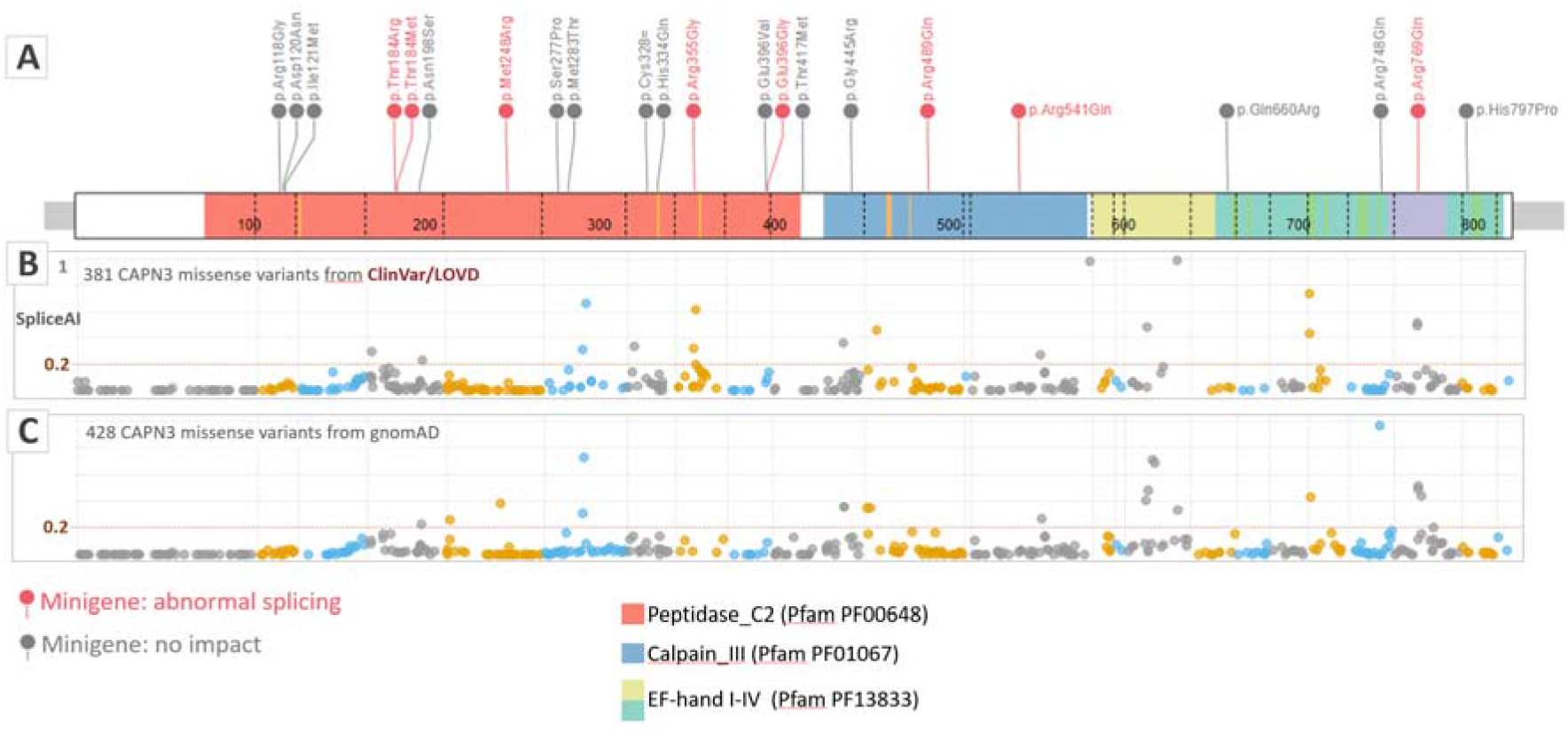
Deep exonic missense variants in *CAPN3* gene. **A**. Twenty one clinically relevant missense variants tested by minigene assay are shown (variants with effect on splicing are in red, variants without impact are in grey). Exon-intron junctions of the *CAPN3* gene as well as the functional domains of calpain 3 are visualized. B. Clinically relevant *CAPN3* variants (pathogenic, likely pathogenic and uncertain/conflicting) from LOVD and ClinVar are visualized along the *CAPN3* gene with SpliceAI scores on the y axis. C. Deep exonic missense variants from gnomADv2.1.1 are visualized along the *CAPN3* gene with SpliceAI scores on the y axis. Variants from adjacent exons are colored differently to visualize the exon junctions. Only variants outside of +/− 2nt from exon borders are shown. Vertical grids in panels B and C correspond to exon boundaries.

**Table 1.** List of 21 deep exonic missense variants tested by functional minigene assay.

| Chr | Start | Ref | Alt | c. | p. | ACMG initial | ACMG post minigene | Minigene | HSF | MaxEntScan | MutPred Splice | SpliceAI | CADD | REVEL |
| --- | --- | --- | --- | --- | --- | --- | --- | --- | --- | --- | --- | --- | --- | --- |
| 15 | 42676723 | A | G | c.352A>G | p.Arg118Gly | likely pathogenic | likely pathogenic | No impact | Alteration of an exonic ESE site. Potential alteration of splicing. | no data | 0.27 | 0.02 | 27.9 | 0.901 |
| 15 | 42676729 | G | A | c.358G>A | p.Asp120Asn | pathogenic | pathogenic | No impact | Alteration of an exonic ESE site. Potential alteration of splicing. | no data | 0.39 | 0.03 | 33 | 0.902 |
| 15 | 42676734 | C | G | c.363C>G | p.Ile121Met | likely pathogenic | likely pathogenic | No impact | Alteration of an exonic ESE site. Potential alteration of splicing. | no data | 0.33 | 0.05 | 27.5 | 0.89 |
| 15 | 42680003 | C | G | c.551C>G | p.Thr184Arg | VUS | likely pathogenic | Abnormal splicing | Creation of an exonic ESS site. Alteration of an exonic ESE site. Potential alteration of splicing. | **impact** | **0.85** | 0.02 | 32 | 0.948 |
| 15 | 42680003 | C | T | c.551C>T | p.Thr184Met | VUS | VUS | Abnormal splicing | Alteration of an exonic ESE site. Potential alteration of splicing. | no data | 0.24 | 0.01 | 33 | 0.87 |
| 15 | 42680045 | A | G | c.593A>G | p.Asn198Ser | likely pathogenic | likely pathogenic | No impact | Activation of an exonic cryptic donor site. Creation of an exonic ESS site. Potential alteration of splicing. | **impact** | **0.83** | **0.23** | 23.4 | 0.769 |
| 15 | 42681236 | T | G | c.743T>G | p.Met248Arg | likely pathogenic | pathogenic | Abnormal splicing | Creation of an exonic ESS site. Alteration of an exonic ESE site. Potential alteration of splicing. | **impact** | **0.81** | 0.05 | 24.7 | 0.851 |
| 15 | 42682178 | T | C | c.829T>C | p.Ser277Pro | VUS | VUS | No impact | Alteration of an exonic ESE site. Potential alteration of splicing. | no data | 0.16 | 0.02 | 16.36 | 0.491 |
| 15 | 42682197 | T | C | c.848T>C | p.Met283Thr | pathogenic | pathogenic | No impact | Alteration of an exonic ESE site. Potential alteration of splicing. | no data | 0.37 | 0.03 | 15.78 | 0.537 |
| 15 | 42684893 | C | G | c.1002C>G | p.His334Gln | pathogenic | pathogenic | No impact | Activation of exonic cryptic donor and acceptor sites, with presence of one or more cryptic branch point(s). Creation of an exonic ESS site. Alteration of an exonic ESE site. Potential alteration of splicing. | **impact** | 0.58 | 0.12 | 25.8 | 0.826 |
| 15 | 42686487 | C | G | c.1063C>G | p.Arg355Gly | likely pathogenic | pathogenic | Abnormal splicing | Creation of an exonic ESS site. Alteration of an exonic ESE site. Potential alteration of splicing. | **impact** | **0.71** | **0.61** | 32 | 0.853 |
| 15 | 42689069 | A | G | c.1187A>G | p.Glu396Gly | VUS | likely pathogenic | Abnormal splicing | Creation of an exonic ESS site. ESE Broken. Most probably affecting splicing. | no data | 0.57 | 0.09 | 32 | 0.925 |
| 15 | 42689069 | A | T | c.1187A>T | p.Glu396Val | likely pathogenic | likely pathogenic | No impact | Creation of an exonic ESS site. Alteration of an exonic ESE site. Potential alteration of splicing. | no data | 0.51 | 0.05 | 33 | 0.897 |
| 15 | 42691746 | C | T | c.1250C>T | p.Thr417Met | pathogenic | pathogenic | No impact | Creation of an exonic ESS site.<br>Alteration of an exonic ESE site.<br>Potential alteration of splicing. | no impact | 0.14 | 0 | 34 | 0.9 |
| 15 | 42691829 | G | A | c.1333G>A | p.Gly445Arg | pathogenic | pathogenic | No impact | Activation of an exonic cryptic acceptor site, with presence of one or more cryptic branch point(s).<br>Alteration of an exonic ESE site.<br>Potential alteration of splicing. | impact | 0.25 | 0.06 | 34 | 0.993 |
| 15 | 42693950 | G | A | c.1466G>A | p.Arg489Gln | pathogenic | pathogenic | Abnormal splicing | Creation of an exonic ESS site.<br>Alteration of an exonic ESE site.<br>Potential alteration of splicing. | no data | 0.12 | 0 | 34 | 0.881 |
| 15 | 42695077 | G | A | c.1622G>A | p.Arg541Gln | pathogenic | pathogenic | Mild impact | Activation of an exonic cryptic acceptor site, with presence of one or more cryptic branch point(s).<br>Creation of an exonic ESS site.<br>Alteration of an exonic ESE site.<br>Potential alteration of splicing. | impact | 0.26 | 0.02 | 34 | 0.908 |
| 15 | 42701565 | A | G | c.1979A>G | p.Gln660Arg | likely pathogenic | likely pathogenic | No impact | Creation of an exonic ESS site.<br>Alteration of an exonic ESE site.<br>Potential alteration of splicing. | no data | 0.3 | 0.02 | 24.9 | 0.792 |
| 15 | 42702844 | G | A | c.2243G>A | p.Arg748Gln | pathogenic | pathogenic | No impact | Creation of an exonic ESS site.<br>Alteration of an exonic ESE site.<br>Potential alteration of splicing. | no data | 0.22 | 0 | 35 | 0.724 |
| 15 | 42703124 | G | A | c.2306G>A | p.Arg769Gln | pathogenic | pathogenic | Mild impact | Activation of an exonic cryptic acceptor site, with presence of one or more cryptic branch point(s).<br>Creation of an exonic ESS site.<br>Potential alteration of splicing. | impact | 0.41 | 0.49 | 35 | 0.947 |
| 15 | 42703494 | A | C | c.2390A>C | p.His797Pro | VUS | VUS | No impact | Creation of an exonic ESS site.<br>Alteration of an exonic ESE site.<br>Potential alteration of splicing. | no impact | 0.19 | 0.01 | 23.4 | 0.528 |

The pathogenicity of 24 functionally tested variants was scored before and after the minigene results according to ACMG criteria (Richards et al. 2015) with the following modifications. PS3 score was assigned if calpain3 protein has been previously reported to be absent on the muscle biopsy Western Blot (PS3_moderate if reduced expression) in at least two patients either homozygous for the variant or compound heterozygous in trans with a confirmed pathogenic variant (PS3_moderate if absent in one patient, PS3_supporting if decreased in one patient). PS3 score was assigned for variants with a minigene result “Impact on splicing”, PS3_supporting for “Mild impact on splicing”. PP2, PP4 and PP5 criteria were not used, consistent with several other recent updates to the original ACMG criteria (Gelb et al. 2018; Lee et al. 2018; Kelly et al. 2018). PP3 score was assigned for variants with REVEL score above 0.7. PP3 was also assigned if two out of three algorithms (HSF, MaxEntScan, SpliceAI) indicated an impact on splicing. PM1 score was assigned for variants affecting highly conserved positions located in the Calpain catalytic domain (PF00648, amino acids 74-417), in Calpain III domain (PF01067, amino acids 436-579) or in EF-hand domains (amino acids 649-683; 692-718; 722-757; 787-821), as visualized by VarMap web tool (Stephenson et al. 2019). PM3 score (*in trans* with a pathogenic variant) was assigned according to SVI Recommendation for in trans Criterion (PM3) - Version 1.0). PP1 score (segregation data) was assigned according to recommendations by the Hearing Loss ClinGen Working group that focused in part on recessive disorders (Oza et al. 2018). The cut-offs for allele frequency criteria were attributed as following: PM2 - 0.02%, BS1 - 0.5%, BA1 - 5%.

Data analysis and visualization was performed with R 3.5.3 using packages dplyr (v.0.8.5, https://dplyr.tidyverse.org/) and ggplot2 (3.2.1, https://ggplot2.tidyverse.org/). ProteinPaint (https://pecan.stjude.cloud/proteinpaint, (Zhou et al. 2016)) was used to visualize variants on the *CAPN3* gene structure in Figure 1.

## RESULTS

### Functional analysis of deep exonic *CAPN3* missense variants using minigene assay

In order to assess the splicing impact of missense variants located further into exons from the exon-intron junctions (deep exonic missense variants), we used minigene assay to functionally test 21 clinically relevant variants homogeneously distributed along the *CAPN3* gene (Figure 1A, Table 1). Eight out of 21 variants tested showed splicing defects in the minigene assay. Of these, five variants induced expression of a shorter transcript lacking one exon (exon skipping), associated with the decrease in the normal transcript (c.551C>G, p.(Thr184Arg); c.551C>T, p.(Thr184Met); c.743T>G, p.(Met248Arg); c.1063C>G, p.(Arg355Gly) and c.1466G>A, p.(Arg489Gln) (Figure 2). In all of these cases, the skipping of the exon was associated with the loss of the reading frame, probably inducing a truncated protein or no protein at all if nonsense mediated decay of the abnormal transcript was induced. Thus, the amount of calpain3 protein present in muscle cells of patients carrying these variants is likely to be lower than normal.

**Figure 2:**
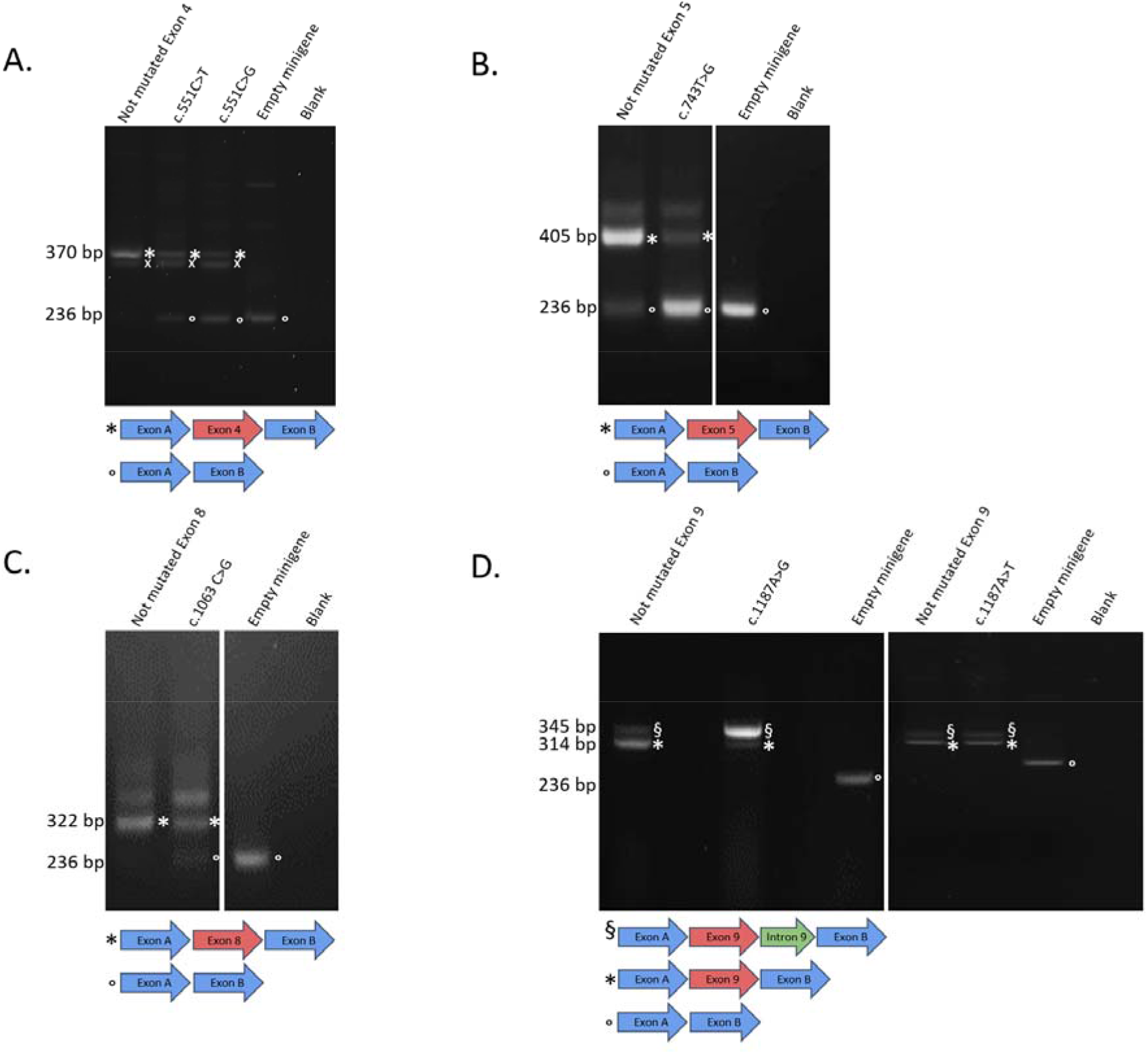
Missense **CAPN3** variants with splicing defects in the minigene assay (exon skipping). RT-PCR gel electrophoresis results from minigene reporter assay, with each band analysed using Sanger sequencing. **A**. Minigene assay for variants c.551 C>G, p.(Thr184Arg) and c.551 C>T, p.(Thr184Met), showing splicing abnormality for both variants. The lower band (indicated with a circle, o) corresponds to the transcript with skipped exon 4 (same as empty minigene). This abnormally spliced transcript is absent from the control lane with the non-mutated construct. The higher band (indicated with an asterisk, *) corresponds to the normal transcript containing exon 4. The normal transcript is present at lower levels for the tested variants compared to control. A non-specific band present in both control and mutated constructs is labeled with an “x”. **B**. Minigene assay for the variant c.743T>G, p.(Met248Arg), showing splicing abnormality. The lower band (indicated with a circle, o) corresponds to the transcript with skipped exon 5 (same as empty minigene). A trace amount of the abnormally spliced transcript is present in the control lane with the non-mutated construct. The higher band (indicated with an asterisk, *) corresponds to the normal transcript containing exon 5. The normal transcript is present at much lower levels for the tested variant compared to control. **C**. Minigene assay for the variant c.1063C>G,p.(Arg355Gly) showing splicing abnormality. The lower band (indicated with a circle, o) corresponds to the transcript with skipped exon 8 (same as empty minigene). This abnormally spliced transcript is absent from the control lane with the non-mutated construct. The higher band (indicated with an asterisk, *) corresponds to the normal transcript containing exon 8. The normal transcript is present at lower levels for the tested variant compared to control. **D**. Minigene assay for the variant c.1466G>A, p.(Arg489Gln) showing splicing abnormality. The lower band (indicated with a circle, o) corresponds to the transcript with skipped exon 11 (same as empty minigene). This abnormally spliced transcript is absent from the control lane with the non-mutated construct. The higher band (indicated with an asterisk, *) corresponds to the normal transcript containing exon 11. The normal transcript is present at lower levels for this variant compared to control.

Minigene assay for the variant c.1187A>G, p.(Glu396Gly) showed presence of a transcript with an abnormal inclusion of the first 31 nucleotides of intron 9 (Figure 3A). The normal transcript was present at a lower level compared to control. Decrease in the normal transcript associated with predicted frameshift in the abnormally spliced transcript will likely lead to lower expression of normal calpain3 protein. Interestingly, minigene assay did not show any splicing abnormality for another nucleotide substitution at the same position (variant c.1187A>T, p.(Glu396Val)).

**Figure 3:**
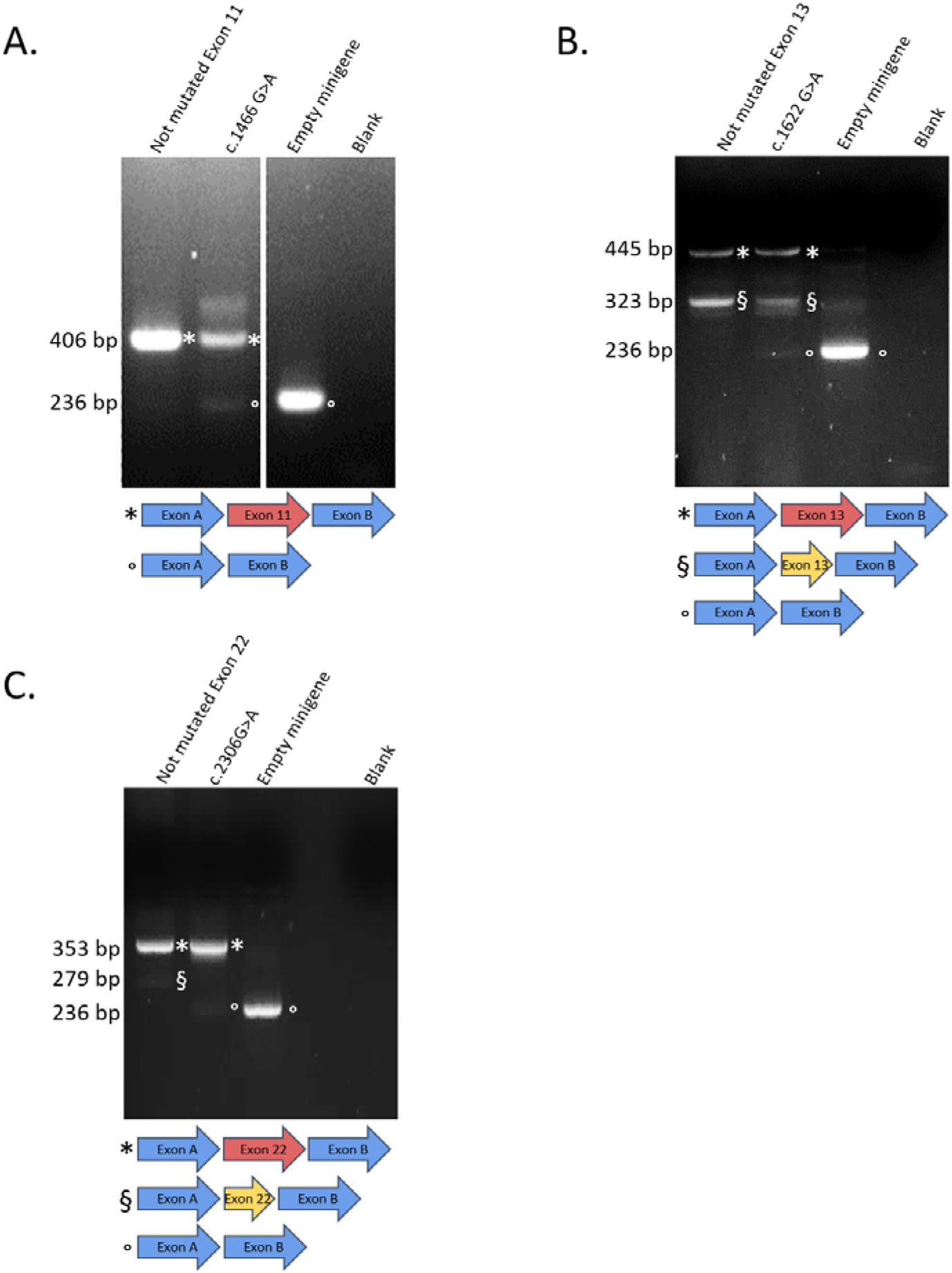
Missense *CAPN3* variants with splicing defects in the minigene assay (other types of splicing abnormalities). RT-PCR gel electrophoresis results from minigene reporter assay, with each band analysed using Sanger sequencing. **A**. Minigene assay for variants c.1187A>G, p.(Glu396Gly) and c.1187A>T, p.(Glu396Val) showing splicing abnormality for c.1187A>G but not for c.1187A>T. The higher band (indicated with §) corresponds to an abnormal transcript containing exon 9 and the first 31 nucleotides of intron 9. This transcript is present at high levels for the construct with c.1187A>G variant. Trace amount of this abnormal transcript is also present in other conditions. The normal transcript (indicated with an asterisk, *) is present at much lower levels for the construct with c.1187A>G variant. There is no difference between transcripts observed for the construct with c.1187A>T variant compared to the control. The empty minigene is indicated with a circle (o). **B**. Minigene assay for the variant c.1622G>A, p.(Arg541Gln) showing splicing patterns different from that of the control construct. The lower band (indicated with a circle, o) corresponds to the transcript with skipped exon 13 (same as empty minigene). This abnormally spliced transcript is absent from the control lane with the non-mutated construct. The higher band (indicated with an asterisk, *) corresponds to the normal transcript containing exon 13. The normal transcript seems to be present at similar levels for mutated as well as for control constructs. A third band (indicated with §) corresponds to the exon 13 missing the last 87 nucleotides (no frameshift). This band is present at high levels in the control and at lower levels in the assay with the mutated construct. **C**. Minigene assay for the variant c.2306G>A, p.(Arg769Gln) showing splicing patterns different from that of the control construct. The lower band (indicated with a circle, o) corresponds to the transcript with skipped exon 22 (same as empty minigene). This abnormally spliced transcript is absent from the control lane with the non-mutated construct. The higher band (indicated with an asterisk, *) corresponds to the normal transcript containing exon 22. The normal transcript seems to be present at similar levels for mutated as well as for control constructs. A third band (indicated with §) corresponds to the transcript containing only the first 43 nucleotides of exon 22. This band is present in the control and absent in the assay with the mutated construct.

Minigene assays for two variants (c.1622G>A, p.(Arg541Gln) and c.2306G>A, p.(Arg769Gln)) showed presence of abnormally spliced transcripts, but the normal transcript expression seemed to be comparable to that of control (FIgure 3B, C). We therefore concluded that the impact of these variants on splicing is likely to be milder compared to the variants described above.

### Localization of deep exonic variant with effect on splicing within exons

In order to explore the localization of splice-affecting exonic missense variants, we calculated distances from these variants to 5’ and 3’ ends of the exons. These variants were located in seven different exons (4, 5, 8, 9, 11, 13 and 22). For exons larger than 100nt, we plotted the splice-affecting exonic variants based on their distance from either 5’ or from 3’ exon ends (Figure 4). For two smaller exons (exons 8 and 9), distances from both ends of the exon are shown (Figure 4). Interestingly, two deep exonic variants with impact on splicing (c.743T>G, p.(Met248Arg) in exon5 and c.1466G>A, p.(Arg489Gln) in exon 11) were located at the exact same distance of 59 nucleotides from the 3’ end of the exon.

**Figure 4:**
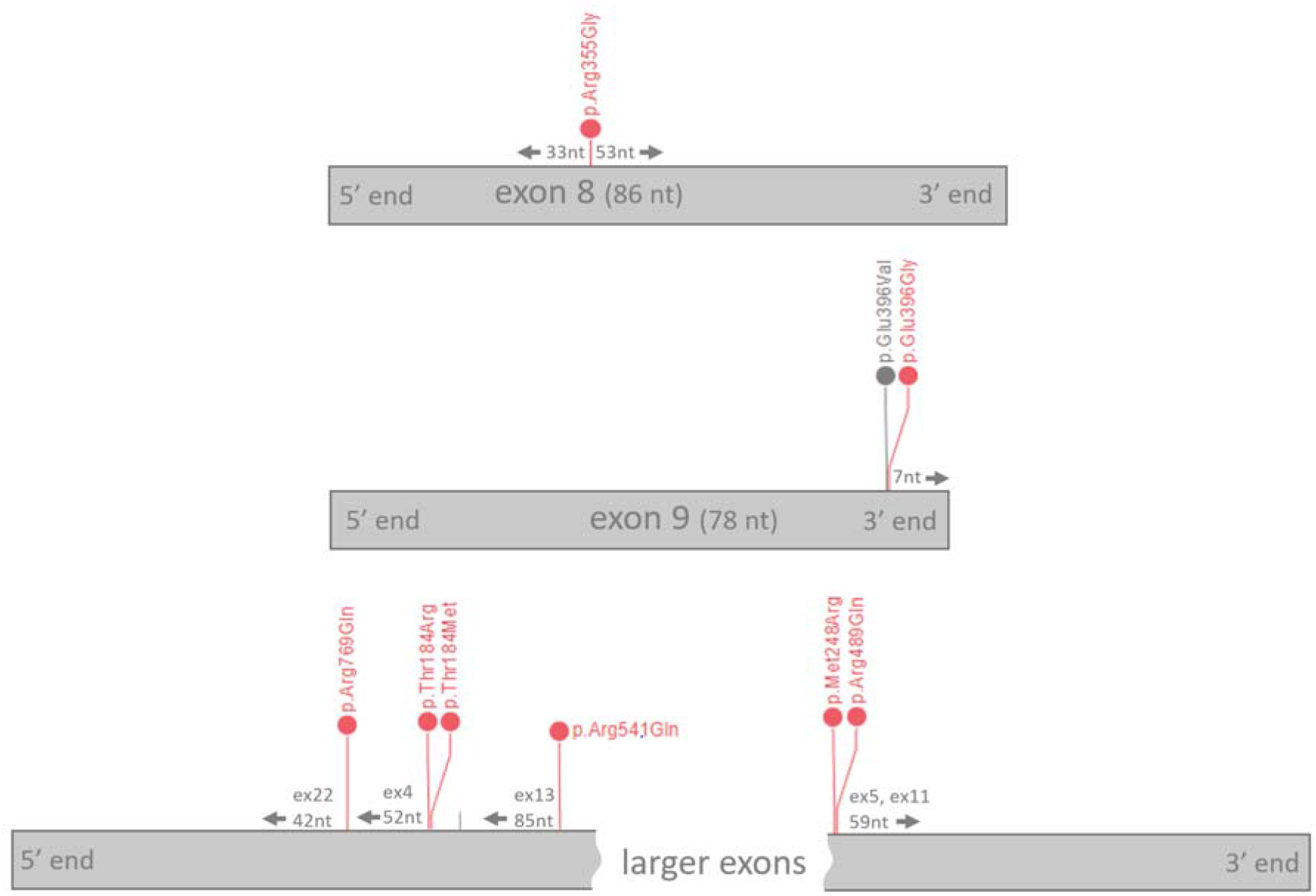
Distance from exonic borders of deep exonic variants with splice impact in minigene assay. Eight missense variants with impact on splicing (red) are visualized along the exons of *CAPN3* gene. Distances from both 5’ and 3’ ends are shown for smaller exons (< 100bp).

We then screened all known *CAPN3* missense variants located in deep exonic regions for potential effect on splicing. To do so, we constituted a set of clinically relevant missense variants (pathogenic, likely pathogenic and uncertain/conflicting) from LOVD and ClinVar databases (Supplementary Figure 1). We then selected only the missense variants located outside of canonical splice sites and used SpliceAI score (Jaganathan et al. 2019) to estimate the putative effect on splicing for each variant. The obtained variants were plotted along the *CAPN3* gene with exon boundaries marked (Figure 1B). Deep exonic missense variants present in the general population (gnomADv2.1.1) were also analyzed (Figure 1C). As seen from these results, certain exonic regions contained clusters of variants with higher SpliceAI scores.

### Splice-prediction algorithm scores do not correlate with the minigene assay results for deep exonic *CAPN3* missense variants

Despite critical need for genetic diagnostics, the currently available splice effect estimation algorithms are still not optimal to correctly predict the impact on splicing, in part due to insufficient number of time-consuming functional validation studies. Our set of *CAPN3* variants functionally tested using the well-established minigene assay provide an excellent opportunity to test the performance of recent splice prediction algorithms. All 21 variants tested in this study were predicted to have an impact on splicing according to HSF, yet only eight showed splicing alteration in the minigene assay, demonstrating a significant false positive rate. The splice effect predictions by MaxEntScan (Yeo and Burge 2004) are shown in Figure 5. Five out of eight variants were correctly predicted as splice-disrupting by MaxEnsScan; for the remaining three variants, no results were available with the matrix used. However, three variants were falsely predicted as splice disrupting. SpliceAI scores for the 21 variants are also shown in Figure 5. Only two out of eight variants thatshowed splicing anomaly on minigene assay had SpliceAI scores above 0.2 (c.1063C>G, p.(Arg355Gly) and c.2306G>A, p.(Arg769Gln)). Of these, c.2306G>A, p.(Arg769Gln) had a mild impact on splicing, since the presence of an abnormally spliced transcript was associated with an unchanged level of normal transcript. Predictions of MutPred Splice, a machine learning-based algorithm designed specifically for exonic variants (Mort et al. 2014), was slightly better than SpliceAI, as it also identified c.551C>G, p.(Thr184Arg) variant (Figure 5). Thus, the majority of the variants with abnormal splicing in minigene assay were not predicted as splice-affecting by SpliceAI and MutPred Splice (6/8 and 5/8 false negatives respectively). One variant (c.593A>G,p.(Asn198Ser)) was predicted as splice disrupting by both algorithms, yet showed no effect on splicing in minigene assay. All eight variants with abnormal splicing in minigene assay had SCAP (Jagadeesh et al. 2019) and MMSplice scores below threshold. We also evaluated predicted impact on protein function, using REVEL and CADD scores (Rentzsch et al. 2019; Ioannidis et al. 2016), (Table 1). All of the eight missense variants with impact on splicing were also predicted to be deleterious at the protein level. Indeed, five variants (p.Thr184Met, p.Thr184Arg, p.Met248Arg, p.Arg355Gly and p.Glu396Gly) affected conserved residues of the Calpain-like peptidase C2 domain (PF00648). Two variants (p.Arg489Gln and p.Arg541Gln) changed conserved amino acids located in the Calpain III domain (PF01067), while p.Arg769Gln affected a conserved amino acid that is expected to participate in protein-protein interaction in a proximity of EF-hand domain (PF13202).

**Figure 5:**
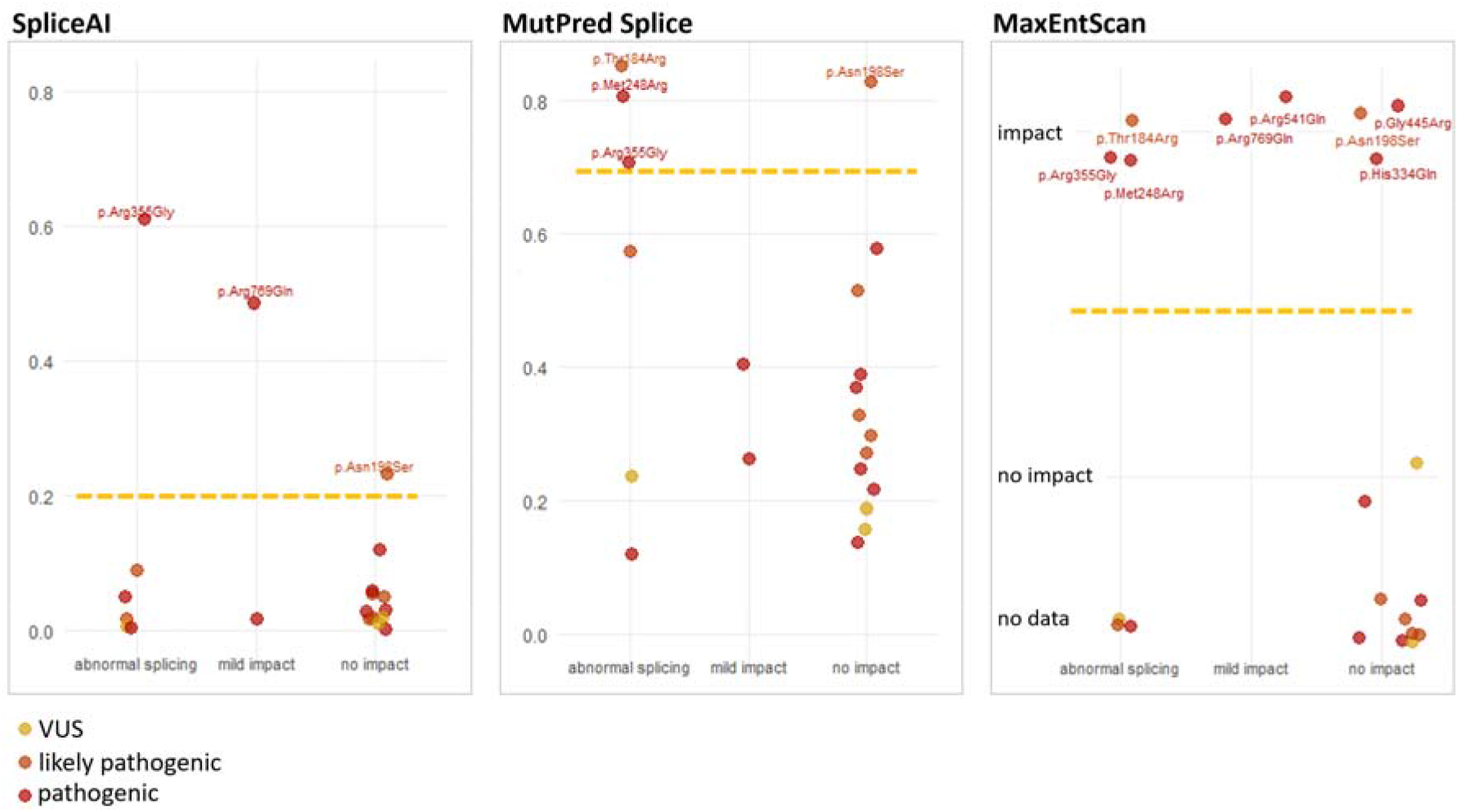
Performance of algorithms predicting impact of splicing for deep exonic variants. Results of three splice prediction computational tools are shown for 21 *CAPN3* missense variants tested by minigene assay. The variants are separated into three groups depending on the minigene result: “Abnormal splicing”, “Mild impact” or “No impact”. The score cut-off above which a variant is predicted to affect splicing is 0.2 for SpliceAI and 0.7 for MutPred Splice.

### Pathogenic classification for seven out of 24 of variants was modified based on the results of the minigene assay

Since most of the variants selected for testing by minigene were identified before the ACMG recommendations for assessing variant pathogenicity, we classified each variant according to the criteria modified for *CAPN3* gene as described in Materials and Methods. The details of evidence used to classify each variant are available in the Supplementary Table 1. The results of the minigene assay, allowing to assign additional PS3 or PS3_sup scores, were then taken into account to calculate “post-minigene ACMG classification” for each variant. For seven out of 24 variants tested by minigene, ACMG classification was modified based on the results of this functional assay.

## DISCUSSION

Despite recent advances in genomic medicine, the pathogenicity of ma variants identified as part of a diagnostic sequencing test is often not well-established. Bioinformatic tools aimed at predicting the identified variant’s effect on protein function or on mRNA splicing therefore play an important role in the diagnostic process (Hu et al. 2019). However, these algorithms do not perform equally well for all types of variants, ultimately requiring a functional confirmation of the pathogenicity prediction. Interpretation of variant pathogenicity is often guided by its location - intronic variants as well as exonic variants in the proximity of exon-intron junctions are usually suspected to disrupt mRNA splicing, while missense variants in the coding regions are considered as potentially altering protein function. Several studies have previously drawn attention to the possibility of exonic variants disrupting normal splicing (Savisaar and Hurst 2017; Kergourlay et al. 2014). In this study, we systematically screened 16 exons of *CAPN3* gene for this type of clinically important variants, by using minigene assay to functionally test missense variants located outside of canonical splice regions (deep exonic variants). We detected splicing abnormalities for eight out of 21 missense variants tested. Interestingly, these eight splice-disrupting missense variants were also predicted to have an impact on protein function due to the amino acid change. It is difficult to disentangle the pathogenic effect on the protein from the effect on mRNA splicing for these variants. It is possible that both mechanisms contribute to the overall pathogenicity in vivo. The interplay between variant’s effect on protein function and its impact on splicing could potentially have important consequences for cell function and disease expression. For example, the effect of dominant negative pathogenic variants on protein level could be downgraded if their splicing effect is increased, reducing the amount of toxic transcript. Recent studies have shown significant variability in mRNA splicing patterns between individuals and human populations in general (Lu, Jiang, and Xing 2012; Park et al. 2018). Thus, one could imagine that the frequency of a confirmed pathogenic dominant negative variant could be higher in a particular population if this variant has greater impact on splicing in this population. On the contrary, if the main pathogenic effect of a variant comes from its impact on splicing, pathogenic abnormally-spliced transcripts could be expressed at lower levels in certain individuals or populations carrying other splice-modifying variants, leading to milder expression of the disease (Cooper et al. 2013). One example of such a variant could be c.551C>T:p.(Thr184Met), one of the variants inducing abnormal splicing in our minigene assay. This missense variant has long been considered as pathogenic, responsible for milder forms of LGMDR1 (Anderson et al. 1998; Richard et al. 1999; Fanin et al. 2004; Barp et al. 2020). This variant had very low frequency in the initially tested subpopulations (0.001% in Europeans, 2 in 129164 alleles in gnomADv2.1.1). However, with more sequencing data now available from different populations, it turns out this variant has 3.3% frequency in African and 0.09% in Latino populations, leading to its reclassification as Benign or Likely benign by many diagnostic laboratories (See Supplementary Table 1 for more details). Given our new evidence of splicing effect for this variant, more studies are needed in order to establish its role in LGMDR1. Another example of a splice-affecting variant with high population frequency (1% in Finnish) is c.1746-20C>G (Nascimbeni et al. 2010). Interestingly, differences in expression of abnormal transcripts was detected between LGMDR1 patients carrying this variant (Nascimbeni et al. 2010), indicating an inter-individual variability of splicing at this locus.

Given the high clinical importance of accurately detecting splice-disrupting variants, a large number of splice effect prediction algorithms have been developed. Earlier tools, such as MaxEntScan and HSF (Yeo and Burge 2004; Desmet et al. 2009), are still widely used in diagnostics, while more recent machine-learning based approaches, such as SpliceAI, S-CAP and MMSplice, are also now gaining significant attention (Rowlands, Baralle, and Ellingford 2019; Jaganathan et al. 2019; Cheng et al. 2019; Jagadeesh et al. 2019). For example, SpliceAI was found to have a positive predictive value of 86% and was selected as the most efficient method for variant prioritization in a recent study from Genomics England Research Consortium (Ellingford et al., 2019, https://doi.org/10.1101/781088). However, these prediction tools were designed and tested using the already known splice variants, most of which are intronic or located at canonical splice sites. In this study, we focused on potential splice-disrupting missense variants located deeper in exons, a group of variants that have been relatively little explored (Savisaar and Hurst 2017). We found the performance of the tools tested suboptimal, as they failed to predict the splicing effect for more than half of the variants showing impact on splicing in the minigene assay. Our results are consistent with a previous study of *BRCA1* and *BRCA2* that found that all variants that affected splicing in a minigene assay while predicted not to disrupt splicing by five different algorithms were located in exons (Théry et al. 2011). Taken together, these findings suggest that exonic variants with effect on splicing might be more difficult to predict using currently available algorithms. The splice-disrupting exonic variants identified in this study could contribute to the new dataset of this type of variants in order to develop more efficient splice-predicting tools.

Exonic elements regulating splicing consist of ESEs (Exonic Splice Enhancers) that promote the splicing of the exon and ESSs (Exonic Splice Silencers) that induce exon exclusion from the mature mRNA (Ohno, Takeda, and Masuda 2018). The density of these elements tends to rise towards the exon-intron junctions (Fairbrother et al. 2004; Woolfe, Mullikin, and Elnitski 2010), thus increasing the probability that a variant present in these regions affects splicing. Interestingly, seven out of eight splice-disrupting missense variants identified in this study were located more than 30 nucleotides away from the exon border. Distribution of regulatory splicing information in the deep exonic regions is less clear. Two models have been previously proposed for functional splice elements in exonic regions - rare regions under strong purifying selection vs. multiple lowly constrained regions (Savisaar and Hurst 2017). Our results are more consistent with the second model, since none of the exonic variants with impact on splicing described in our study are located in regions of high constraint (CCR > 90 (Havrilla et al. 2019)). Indeed, only a small region of exon 5 of *CAPN3* is constrained using this method, most likely due to the fact that the vast majority of *CAPN3* pathogenic variants are responsible for recessive disease.

Previous studies using minigene assays have estimated that about 30% of exonic variants affect splicing (Savisaar and Hurst 2017). Our results are consistent with these findings (eight splice-affecting variants out of 21 tested by minigene), even though all 21 variants tested were predicted to be splice-modifying by HSF, thus overestimating the proportion of splice-affecting exonic variants.

Accessibility and wide use of high throughput sequencing for limb-girdle myopathy diagnosis has led to identification of multiple variants of uncertain pathogenicity, increasing the demand for efficient functional studies. For LGMDR1, presence of calpain3 protein on Western Blot or screening for abnormal transcripts in muscle RNA has proven to be useful to achieve accurate molecular diagnosis. However, both of these approaches require access to patient’s muscle biopsy material. A useful *in vitro* alternative is the well-established minigene assay. It has been previously shown that the results of minigene assays are concordant with the analysis of patient RNA (Tournier et al. 2008; Théry et al. 2011; Bonnet et al. 2008). Here, we developed the minigene approach for the majority of *CAPN3* exons and used it to test 24 variants. The results of the minigene assay allowed assigning additional evidence points according to ACMG recommendations (Richards et al. 2015) leading to change in ACMG classification for seven variants. These results demonstrate the utility of minigene functional assay for diagnostics of calpainopathy and reinforce the need to incorporate functional studies into diagnostic process on a larger scale. The functional assay described here can be directly used by the diagnostic laboratories to test *CAPN3* variants with suspected splicing effect, allowing to establish long-awaited diagnosis for many LGMDR1 patients.

## Acknowledgements

We thank patients and their relatives for their participation in this study. We also acknowledge Alexandra Martins and Christiane Duponchel for generously providing the pCAS2 vector.

---

**Figure S1:**
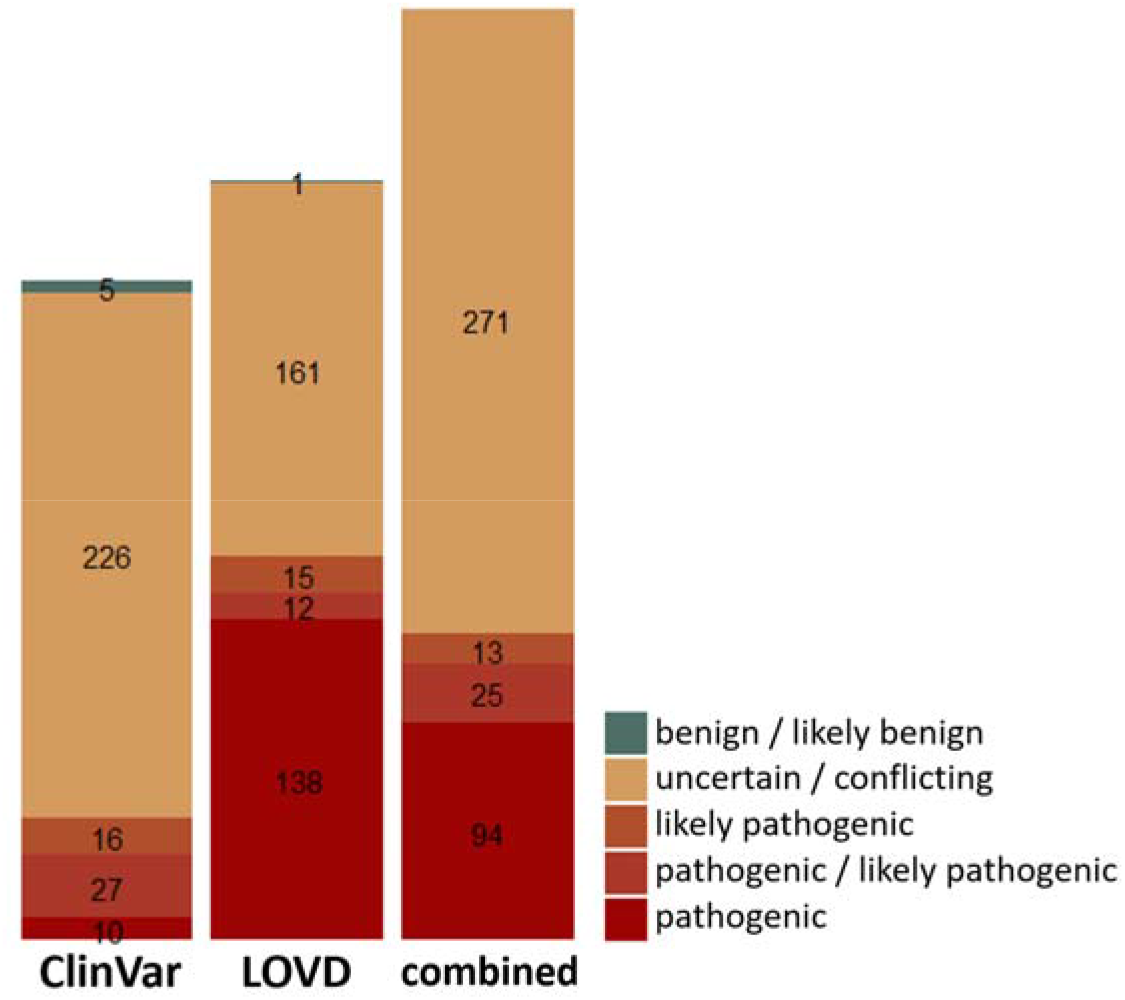
Clinically significant missense variants in *CAPN3* gene. A set of 403 unique clinically significant missense variants was obtained by combining variants from LOVD and ClinVar annotated as non benign in at least one of these databases (327 variants from LOVD, 284 variants from ClinVar). *CAPN3* missense variants from LOVD were assigned the “Clinical Significance” values based on the classification system used in ClinVar (uncertain/conflicting, likely pathogenic, pathogenic/likely pathogenic or pathogenic).

**Table S1:** Description of variants selected to be tested by minigene assay.

| Evidence for each variant | ACMG score before Minigene | ACMG score after Minigene |
| --- | --- | --- |
| CAPN3 NM_000070.2:c.352A>G:p.(Arg118Gly) (hg19)chr15:42676723 Canonical Allele Identifier: CA391997392<br>This variant has previously been reported in a compound heterozygous state in one LGMDR1 patients (PMID:10330340, LOVD Individual ID 00214096), found *in trans* with a pathogenic variant (c.1981_1984del, p.(Ile661Glnfs*20))(**PM3_sup**). It is present in only one individual in gnomADv3 (0.005% 1/18394 in East Asian population) and absent from gnomADv2.1.1 (**PM2**). This variant is located at a conserved residue of the Calpain catalytic domain (PF00648) (**PM1**) and is predicted to affect protein function (CADD 27.9, REVEL 0.901) (**PP3**). Purified rat m-calpain with the corresponding mutation had only 5% of wild-type activity (PMID: 11371436) (**PS3_sup**).<br>HSF: Alteration of an exonic ESE site. Potential alteration of splicing. MaxEntScan: no result. SpliceAI: No impact (0.02)<br>Minigene: “No impact on splicing” | PS3_sup, PM1, PM2, PM3_sup, PP3<br>**Likely pathogenic** | No change |
| CAPN3 NM_000070.2:c.358G>A:p.(Asp120Asn) (hg19)chr15:42676729 Canonical Allele Identifier: CA7510925<br>This variant has previously been reported in a homozygous state in two LGMDR1 patients (PMID: 15689361, LOVD Individual ID 00213833) associated with a decrease of Calpain3 protein on Western Blot. We have identified this variant in a patient from the Timone Calpainopathy cohort, *in trans* with a pathogenic variant, associated with a decrease of Calpain3 on Western Blot (we submitted this variant to LOVD, Individual ID 289314) (**PM3_strong, PS3_mod**). This variant is present at the MAX population frequency of 0.006% (2/30616 of South Asian population, gnomADv2.1.1) (**PM2**). This variant is located at a conserved residue of the Calpain catalytic domain (PF00648) (**PM1**) and is predicted to affect protein function (CADD 33, REVEL 0.902) (**PP3**).<br>HSF: Alteration of an exonic ESE site. Potential alteration of splicing. MaxEntScan: no result. SpliceAI: No impact (0.03)<br>Minigene: “No impact on splicing” | PS3_mod, PM1, PM2, PM3_strong, PP3<br>**Pathogenic** | No change |
| CAPN3 NM_000070.2:c.363C>G:p.(Ile121Met) (hg19)chr15: 42676734 Canonical Allele Identifier: CA269832799<br>There are 3 observations listed for this variant in ClinVar (RCV000595690.1), classified as VUS. No phenotypic information is available for these individuals. There are four additional VUS entries in LOVD (Individual IDs 213632, 222054, 222213, 219478). Each of these individuals carries a known pathogenic CAPN3 variant (one c.2362_2363delinsTCATCT and three c.550del), but no phenotypic information is available.<br>We have identified this variant in a patient from the Timone Calpainopathy cohort, confirmed *in trans* with a pathogenic variant, without Calpain3 decrease on Western Blot (we submitted this variant to LOVD, Individual ID 289315) (**PM3_strong**). This variant is present at the MAX population frequency of 0.01% (1/8718 in African population, gnomADv2.1.1) (**PM2**). This variant is located at a conserved residue of the Calpain catalytic domain (PF00648) (**PM1**) and is predicted to affect protein function (CADD 27.5, REVEL 0.89) (**PP3**).<br>HSF: Alteration of an exonic ESE site. Potential alteration of splicing. MaxEntScan: no result. SpliceAI: No impact (0.05)<br>Minigene: “No impact on splicing” | PM1, PM2, PM3_strong, PP3<br>**Likely pathogenic** | No change |
| CAPN3 NM_000070.2:c. c.498G>A p.Gln166= (hg19)chr15: 42678483 Canonical Allele Identifier: CA7511028<br>This synonymous variant is listed as benign in one individual (PMID10330340, LOVD Individual ID 214423). It is present in only one allele in gnomAD (v2.1.1 and v3, POPMAX 0.005%, 1/18392 East Asian population) (**PM2**). This variant is located at the last nucleotide of exon3 (canonical splice-site). HSF: Alteration of an exonic ESE site. Potential alteration of splicing. MaxEntScan: impact on splicing. SpliceAI: Impact on splicing (DS_DG=0.49, DS_DL=0.77) (**PP3**)<br>Minigene: “Impact on splicing” (**PS3**)<br>Presence of an abnormal transcript with inclusion of first 5bp of intron 3. A second abnormal transcript with exon skipping is also present. Both cause frameshift. No normal transcript is present. | PM2, PP3<br>**VUS** | PS3, PM2, PP3<br>**Likely pathogenic** |
| CAPN3 NM_000070.2:c.551C>T:p.(Thr184Met) (hg19)chr15:42680003 Canonical Allele Identifier: CA152139<br>This variant has been reported in the homozygous state or *in trans* with a pathogenic or likely pathogenic variant in 5 patients with a mild decrease of calpain3 on Western Blot in one case (PMID: 9777948, 10330340, 15221789, 31555977, LOVD:214085, 214156) (**PM3_strong**). This variant has also been published or reported in a database as compound heterozygous with variants that today can be re-classified as VUS or likely benign in 4 individuals (PMID:18055493, 19835634, LOVD individual IDs: 214602, 214606, 214666, 214738). There are five ClinVar entries classifying this variant as Benign or Likely benign based on its frequency in one sub-population (VCF000128572.2). This variant is present at 3.3% in African population (1397/42024, 25 homozygotes gnomADv.3) (**BS1**). This variant is located at a conserved residue of the Calpain catalytic domain (PF00648) (**PM1**) and is predicted to affect protein function (CADD 33, REVEL 0.87) (**PP3**). HSF: Alteration of an exonic ESE site. Potential alteration of splicing. MaxEntScan: no result. SpliceAI: No impact (0.01)<br>Minigene: “Impact on splicing” (**PS3**)<br>Presence of an abnormal transcript lacking exon 4 (causes frameshift). The normal transcript (including exon 4) is expressed at lower level comparing to control. | PM1, PM3_strong, PP3, BS1<br>**VUS** | **PS3**, PM1, PM3, PP3, BS1<br>No change |
| CAPN3 NM_000070.2:c.551C>G:p.(Thr184Arg) (hg19)chr15:42680003 Canonical Allele Identifier: CA391997854<br>This variant has neither been reported in the literature nor submitted to LOVD or ClinVar. It was identified by our laboratory in two siblings affected with LGMD without Western Blot data available, associated to a VUS c.2380+6T>C. We submitted this variant to LOVD (Individual LOVD ID: 289317). This variant is absent from population databases (gnomADv2.1.1 and v3) (**PM2**). This variant is located at a conserved residue of the Calpain catalytic domain (PF00648) (**PM1**) and is predicted to affect protein function (CADD 32, REVEL 0.95) (**PP3**).<br>HSF: Creation of an exonic ESS site. Alteration of an exonic ESE site. Potential alteration of splicing. MaxEntScan: impact on splicing. SpliceAI: No impact (0.02) (**PP3**)<br>Minigene: “Impact on splicing” (**PS3**)<br>Presence of an abnormal transcript lacking exon 4 (causes frameshift). The normal transcript (including exon 4) is expressed at lower level | PM1, PM2, PP3<br>**VUS** | **PS3**, PM1, PM2, PP3<br>**Likely pathogenic** |
| comparing to control. |  |  |
| CAPN3 NM_000070.2:c.593A>G:p.(Asn198Ser) (hg19)chr15:42680045 Canonical Allele Identifier: CA7511066<br>This variant has previously been reported in a compound heterozygous state in one LGMDR1 patients (PMID:16650086, LOVD Individual ID 00214395, *in trans* with NM_000070.2:c.146G>A:p.Arg49His) (**PM3_Sup**) associated with an absence of Calpain3 protein on Western Blot (**PS3_mod**). There are two additional entries in LOVD classified as VUS (LOVD Individual IDs 215020, 221029). ClinVar contains three entries for this variant classified as VUS in one affected individual and two individuals without phenotypic information (SCV000645506, SCV000800198, SCV000568284). This variant is present at the MAX population frequency of 0.046% (1/2154 alleles of "Other" population in gnomADv.3), but frequencies in other populations are 10x lower. Given the relatively low number of alleles in "Other" population as well as lack of additional information about the origins of individuals in this group, **PM2** was assigned for this variant. This variant is located at a conserved residue of the Calpain catalytic domain (PF00648) (**PM1**) and is predicted to affect protein function (CADD 23.4, REVEL 0.769) (**PP3**).<br>HSF: Activation of an exonic cryptic donor site. Creation of an exonic ESS site. Potential alteration of splicing. MaxEntScan: impact on splicing. SpliceAI : Abnormal splicing (0.23) (**PP3**)<br>Minigene: "No impact on splicing" | PM1, PM2, PS3_mod, PM3_sup, PP3<br>**Likely pathogenic** | No change |
| CAPN3 NM_000070.2:c.743T>G:p.(Met248Arg) (hg19)chr15:42681236 Canonical Allele Identifier: CA7511100<br>This variant has previously been reported in a compound heterozygous state in one LGMDR1 patient (*in trans* with NM_000070.2:c.2306G>A) as well as in 4 homozygous individuals from one family (PMIDs:16650086, 12461690) (**PM3**). Western Blot showed reduction of Calpain3 in all of these cases (**PS3_mod**). There is one entry in LOVD for this variant, classified as VUS (LOVD Individual ID 214004). This variant is present at the MAX population frequency of 0.007% (1/13646 alleles from Latino population in gnomADv2.1.1) (**PM2**). This variant is located in the Calpain catalytic domain (PF00648), but the amino acid residue not conserved. It is predicted to affect protein function (CADD 24.7, REVEL 0.851) (**PP3**).<br>HSF: Creation of an exonic ESS site. Alteration of an exonic ESE site. Potential alteration of splicing. MaxEntScan: impact on splicing. SpliceAI: No impact (0.05)(**PP3**)<br>Minigene: "Impact on splicing" (**PS3**)<br>Presence of an abnormal transcript lacking exon 5 (causes frameshift). The normal transcript (including exon 5) is expressed at much lower level comparing to control. | PM2, PS3_mod, PM3, PP3<br>**Likely pathogenic** | PM2, **PS3**, PS3_mod, PM3, PP3<br>**Pathogenic** |
| CAPN3 NM_000070.2:c.829T>C:p.(Ser277Pro) (hg19)chr15:42682178 Canonical Allele Identifier: CA391998485<br>This variant has neither been reported in the literature nor submitted to LOVD or ClinVar. It was identified by our laboratory in one patient affected with LGMDR1 with a decrease of calpain3 on Western Blot, associated to a VUS c.1746-20C>G (**PS3_mod**). We submitted this variant to LOVD (Individual LOVD ID: 289316). This variant is absent from population databases (gnomADv2.1.1 and v3) (**PM2**). This variant is located in the Calpain catalytic domain (PF00648), but the amino acid residue not conserved. It is not predicted to affect protein function (CADD 16.36, REVEL 0.491).<br>HSF: Alteration of an exonic ESE site. Potential alteration of splicing. MaxEntScan: no result. SpliceAI: No impact (0.02) | PS3_mod, PM2<br>**VUS** | No change |
| Minigene: "No impact on splicing" |  |  |
| CAPN3 NM_000070.2:c.848T>C:p.(Met283Thr) (hg19)chr15:42682197 Canonical Allele Identifier: CA391998519<br>This variant has previously been reported in the compound heterozygous state in three patients and in the homozygous state in one LGMDR1 patient (PMIDs 15221789, 25046369, 26404900, LOVD Individual ID 213845). The homozygous patient had no Calpain3 protein on Western Blot (PMID: 15221789). This variant was identified by our laboratory in a patient affected with LGMDR1, *in trans* with a pathogenic variant, associated with an absence of Calpain3 on Western Blot (we submitted this variant to LOVD, Individual ID 289318) (**PS3, PM3_strong**). There is one submission in ClinVar for this variant, classified as Likely Pathogenic (SCV000796489), bringing the total number of reported patients to six. This variant is absent from population databases (gnomADv2.1.1 and v3) (**PM2**). This position is located in the Calpain catalytic domain (PF00648), but it is not conserved and the variant is not predicted to affect protein function (CADD 15.78, REVEL 0.537). HSF: Alteration of an exonic ESE site. Potential alteration of splicing. MaxEntScan: no result. SpliceAI: No impact (0.03)<br>Minigene: "No impact on splicing" | PS3, PM2, PM3_strong<br>**Pathogenic** | No change |
| CAPN3 NM_000070.2:c.984C>T:p.(Cys328=) (hg19)chr15:42684875 Canonical Allele Identifier: CA179838<br>This synonymous variant is present at 1% in European population (1327/129002, 12 homozygotes gnomADv.3) (**BS1**). There four entries in ClinVar classified Benign or Likely Benign (VCV000166787.2)<br>HSF: Activation of an exonic cryptic donor site. Alteration of an exonic ESE site. Potential alteration of splicing. MaxEntScan: no result. SpliceAI: No impact (0.07)<br>Minigene: "No impact on splicing" (**BS3**) | BS1<br>**Likely Benign** | BA1, **BS3**<br>**Benign** |
| CAPN3 NM_000070.2:c.1002C>G:p.(His334Gln) (hg19)chr15:42684893 Canonical Allele Identifier: CA391998854<br>This variant has previously been reported in the compound heterozygous state in three LGMDR1 patients from one family (PMID: 9150160, LOVD individual IDs 213909, 214972) (**PM3, PP1_strong**). Other variants affecting the same amino acid (p.His334Tyr and p.His334Leu) have been reported in LGMDR1 patients in the literature and ClinVar, (PMIDs: 16141003, 18055493, VCV000285580.3) (PM5 not attributed since PM1 is also present). The affected amino acid corresponds to the conserved catalytic residue of Calpain catalytic domain (PF00648) (**PM1**). This variant is predicted to affect protein function (CADD 25.8, REVEL 0.826) (**PP3**). This variant is present at the MAX population frequency of 0.0009% (1/113562 alleles from European population in gnomADv2.1.1) and absent from all other populations (**PM2**)<br>HSF: Activation of exonic cryptic donor and acceptor sites, with presence of one or more cryptic branch point(s). Creation of an exonic ESS site. Alteration of an exonic ESE site. Potential alteration of splicing. MaxEntScan: impact on splicing. SpliceAI: No impact (0.12) (**PP3**)<br>Minigene: "No impact on splicing" | PM1, PM2, PM3, PP1_strong, PP3<br>**Pathogenic** | No change |
| CAPN3 NM_000070.2:c.1063C>G:p.(Arg355Gly) (hg19)chr15:42686487 Canonical Allele Identifier: CA391998989 | PM1, PM2, PM3_sup, | **PS3**, PM1, PM2, |
| This variant has previously been reported in the compound heterozygous state in one LGMDR1 patient (PMID:16650086, LOVD individual ID 214380) **(PM3_sup)**. A known pathogenic variant affecting the same amino acid (p.Arg355Trp) has been reported in LGMDR1 patients in the literature and ClinVar (PMIDs 26404900, 23821418, 18854869, 19364062, 18073330, 17994539, 17236769, 15884399, 15689361, 29970176, VCV000281254.2) ) (PM5 not attributed since PM1 is also present). It is absent from general population (gnomADv2.1.1, v3) **(PM2)**. This variant is located at a conserved residue of the Calpain catalytic domain (PF00648) **(PM1)** and is predicted to affect protein function (CADD 32, REVEL 0.853) **(PP3)**.<br>HSF: Creation of an exonic ESS site. Alteration of an exonic ESE site. Potential alteration of splicing. MaxEntScan: impact on splicing. SpiceAI: Abnormal splicing (DS_AL=0.61) ( **(PP3)**)<br>Minigene: “Impact on splicing” **(PS3)**<br>Presence of an abnormal transcript lacking exon 8 (causes frameshift). The normal transcript (including exon 8) is expressed at lower level compared to control. | PP3<br>**Likely pathogenic** | PM3_sup, PP3<br>**Pathogenic** |
| CAPN3 NM_000070.2:c.1187A>G:p.(Glu396Gly) (hg19)chr15:42689069 Canonical Allele Identifier: CA7511268<br>There are two entries in LOVD and ClinVar for this variant classified at VUS (VCV000497565.1, LOVD Individual ID 222031). It is present at the MAX population frequency of 0.003% (2/64564 alleles from European population in gnomADv3)**(PM2)**. This variant is located at a conserved residue of the Calpain catalytic domain (PF00648) **(PM1)** and is predicted to affect protein function (CADD 32, REVEL 0.925) **(PP3)**.<br>HSF: Creation of an exonic ESS site. ESE Broken. Most probably affecting splicing. MaxEntScan: no result. SpiceAI: No impact (0.09)<br>Minigene: “Impact on splicing” **(PS3)**<br>Presence of an abnormal transcript with inclusion of first 31bp of intron 9 (causes frameshift). The normal transcript is expressed at lower level compared to control. | PM1, PM2, PP3<br>**VUS** | **PS3, PM1, PM2, PP3**<br>**Likely pathogenic** |
| CAPN3 NM_000070.2:c.1187A>T:p.(Glu396Val) (hg19)chr15:42689069 Canonical Allele Identifier: CA391999275<br>This variant has not been reported before. It was identified in our center in one LGMD patient *in trans* with a known pathogenic variant (c.946-1G>A)**(PM3)**. We submitted this variant to LOVD (LOVD Individual ID: 289319). There are two entries in LOVD and ClinVar for another variant affecting the same amino acid (p.Glu396Gly) classified as VUS (VCV000497565.1, LOVD Individual ID 222031). This variant is absent from general population (gnomADv2.1.1, v3) **(PM2)**. It is located at a conserved residue of the Calpain catalytic domain (PF00648) **(PM1)** and is predicted to affect protein function (CADD 33, REVEL 0.897) **(PP3)**.<br>HSF: Creation of an exonic ESS site. Alteration of an exonic ESE site. Potential alteration of splicing. MaxEntScan: no result. SpiceAI: No impact (0.05)<br>Minigene: “No impact on splicing” | PM1, PM2, PM3, PP3<br>**Likely pathogenic** | No change |
| CAPN3 NM_000070.2:c.1250C>T:p.(Thr417Met) (hg19)chr15:42691746 Canonical Allele Identifier: CA7511300<br>This variant has been previously identified in a homozygous or compound heterozygous state in five LGMDR1 patients (PMIDs: 25135358, 25079074, 17702496, 16650086). There are also more than ten LOVD and ClinVar entries for this variant with Pathogenic or Likely Pathogenic | PM3_VS, PM2, PP3, PS3_sup | No change |
| classification (**PM3_VS**). This variant is present at the MAX population frequency of 0.009% (12/129150 alleles from European population in gnomADv2.1.1)(**PM2**). It is predicted to affect protein function (CADD 34, REVEL 0.9) (**PP3**). Functional studies using recombinant rat calpain 2 with the corresponding variant showed a decrease in enzyme activity (PMID: 19226146) (**PS3_sup**).<br>HSF: Creation of an exonic ESS site. Alteration of an exonic ESE site. Potential alteration of splicing. MaxEntScan: no impact on splicing. SpliceAI: No impact (0.00)<br>Minigene: “No impact on splicing” | **Pathogenic** |  |
| CAPN3 NM_000070.2:c.1333G>A:p.(Gly445Arg) (hg19)chr15:42691829 Canonical Allele Identifier: CA7511320<br>This variant has been reported in multiple LGMDR1 individuals in a heterozygous state. A manuscript providing evidence that this variant is responsible for a dominant form of the calpainopathy is currently in revision (Cerino et al.)<br>The classification provided here is for the AR mode of inheritance for this variant.<br>This variant has been reported in two patients in compound heterozygous state with a likely pathogenic variant (PMIDs: 10330340, 17994539, LOVD Individual IDs: 214106, 214546), with reduced or absent protein on Western Blot. One additional homozygous patient with the absence of calpain3 on Western Blot is reported in LOVD (Individual ID 213752). There are four Pathogenic or Likely Pathogenic ClinVar entries for this variant (SCV000791979, SCV000337170, SCV000953639, SCV001164533), including one LGMDR1 patient with a pathogenic variant *in trans* (**PM3_strong, PS3**). There is an LOVD entry for a different nucleotide variant (c.1333G>C) leading to the same amino acid change, *in trans* with a pathogenic variant (Individual ID 215007) and a ClinVar entry classified as Likely pathogenic (VCV000596306).<br>This variant is present at the MAX population frequency of 0.002% (3/107778 alleles from European population in gnomADv2.1.1)(**PM2**). It is located in Calpain III domain (PF01067) and is affecting a conserved amino acid (**PM1**) and is predicted to affect protein function (CADD 34, REVEL 0.99) (**PP3**).<br>HSF : Activation of an exonic cryptic acceptor site, with presence of one or more cryptic branch point(s). Alteration of an exonic ESE site. Potential alteration of splicing. MaxEntScan: impact on splicing. SpliceAI: No impact (0.06) (**PP3**)<br>Minigene: “No impact on splicing” | PM3_strong, PS3, PM1, PM2, PP3<br>**Pathogenic** | No change |
| CAPN3 NM_000070.2:c.1466G>A:p.(Arg489Gln) (hg19)chr15:42693950 Canonical Allele Identifier: CA7511363<br>This variant has been reported in more than six patients in a compound heterozygous state with a confirmed pathogenic variant (PMIDs: 1033034, 14578192, 17994539, 30564623, LOVD Individual IDs: 214142, 214297, 215059) (**PM3_VS**). Calpain3 protein was reduced in Western Blot in one case (**PS3_sup**). There are four Pathogenic/Likely Pathogenic entries for this variant in ClinVar (SCV000798519, SCV000344033, SCV000616671, SCV000612633). This variant is present at the MAX population frequency of 0.048% (20/42020 of African population, gnomADv3). It is located in Calpain III domain (PF01067) and is affecting a conserved amino acid that is expected to participate in protein-protein interaction (**PM1**). It is predicted to affect protein function (CADD 34, REVEL 0,881) (**PP3**). This variant has also been shown to reduce the autocatalytic activity of calpain3 protein (PMID: 14578192) (**PS3**).<br>HSF: Creation of an exonic ESS site. Alteration of an exonic ESE site. Potential alteration of splicing. MaxEntScan: no result. SpliceAI: No impact (0.00) | PM3_VS, PS3, PM1, PP3<br>**Pathogenic** | PM3_VS, PS3, PM1, PP3<br>No change |
| Minigene: “Impact on splicing” (**PS3**)<br>Presence of an abnormal transcript lacking exon 11 (causes frameshift). The normal transcript (including exon 11) is expressed at lower level compared to control. |  |  |
| CAPN3 NM_000070.2:c.1622G>A:p.(Arg541Gln) (hg19)chr15:42695077 Canonical Allele Identifier: CA220342<br>More than ten LGMDR1 patients carrying this variant in compound heterozygous state have been reported in the literature (PMID: 10330340, 16141003, 16411092, 30564623) (**PM3_PVS**). Seven additional individuals with this variant are listed in ClinVar (VCV000092407.3) and LOVD (Individual IDs 214986, 213698). A different substitution of the same amino acid (p.Arg541Trp) is also a known pathogenic variant (PMID: 16141003, 14981715) ) (PM5 not attributed since PM1 is also present). This variant is present at the MAX population frequency of 0.005% (1/18390 of East Asian population, gnomADv2.1.1) (**PM2**). It is affecting a conserved amino acid located in the Calpain III domain (PF01067) (**PM1**) and is predicted to affect protein function (CADD 34, REVEL 0.908) (**PP3**).<br>HSF: Activation of an exonic cryptic acceptor site, with presence of one or more cryptic branch point(s). Creation of an exonic ESS site. Alteration of an exonic ESE site. Potential alteration of splicing. MaxEntScan: impact on splicing. SpiceAI: No impact (0.02) (**PP3**)<br>Minigene: “Mild impact on splicing” (**PS3_sup**)<br>Presence of an abnormal transcript lacking exon 13 (causes frameshift). The normal transcript (including exon 13) is expressed at similar level compared to control. Another abnormal transcript missing the last 87 nucleotides of exon 13 is present for both control and mutated constructs. | PM3_VS, PM2, PP3<br>**Pathogenic** | No change |
| CAPN3 NM_000070.2:c.1913A>C:p.(Gln638Pro) (hg19)chr15:42700521 Canonical Allele Identifier: CA392000906<br>This variant has been previously identified in a compound heterozygous state in one LGMDR1 patient *in trans* with a known pathogenic variant, associated with a decrease of calpain3 on Western Blot (PMID: 10330340, LOVD Individual ID: 214080 (**PM3, PS3_sup**). This variant is absent from population databases (gnomADv2.1.1, v3) (**PM2**). It is not predicted to affect protein function (CADD 16.22, REVEL 0.481). Even though it is located 2nt upstream of the 3’ end of exon 16, HSF prediction was: No significant splicing motif alteration detected. This mutation has probably no impact on splicing. MaxEntScan: no result. SpiceAI: Abnormal splicing (DS_DL=0.72)<br>Minigene: “Impact on splicing” (**PS3**)<br>Presence of an abnormal transcript lacking exon 16 (does not cause frameshift). | PM2, PM3, PS3_sup<br>**VUS** | PM2, PM3, PS3_sup, **PS3**<br>**Likely pathogenic** |
| CAPN3 NM_000070.2:c.1979A>G:p.(Gln660Arg) (hg19)chr15:42701565 Canonical Allele Identifier: CA392001277<br>This variant has previously been reported in a compound heterozygous state in four LGMDR1 patients (PMID: 16650086, 15689361), (**PM3_strong**). Western blot showed complete absence of Calpain3 in one case and trace amounts in two other cases (PMID: 16650086, 15689361), (**PS3**). There is one ClinVar submission for this variant (VCV000558137.1), classified as VUS in May 2018. This variant is absent from population databases (gnomADv2.1.1, v3) (**PM2**). It is predicted to affect protein function (CADD 24.9, REVEL 0.792) (**PP3**).<br>HSF: Creation of an exonic ESS site. Alteration of an exonic ESE site. Potential alteration of splicing. MaxEntScan: no result. SpiceAI: No impact (0.02) | PM3_strong, PS3, PM2, PP3<br>**Pathogenic** | No change |
| Minigene: "No impact on splicing" |  |  |
| CAPN3 NM_000070.2:c.2243G>A:p.(Arg748Gln) (hg19)chr15:42702844 Canonical Allele Identifier: CA345536<br>This variant has been reported in the homozygous or compound heterozygous state in many individuals affected with LGMDR1 (PMID: 12461690, 9777948, 18854869, 27020652, 18563459, 15689361, 19015733, 22443334, 10330340, 26404900, 9762961) (**PM3_VS**). This variant was shown to co-segregate with LGMD in one affected family, with two affected individuals (PMID: 9150160) (**PP1_mod**). There are nine entries in ClinVar classifying this variant as "Pathogenic" or "Likely pathogenic" (VCF000128570). This variant is present at the MAX population frequency of 0.014% (5/35372 of Latino population, gnomADv2.1.1) (**PM2**) and is predicted to affect protein function (CADD 35, REVEL 0.724) (**PP3**). Functional studies using recombinant rat calpain 2 with the corresponding mutation showed a decrease in enzyme activity (PMID: 19226146) (**PS3_sup**).<br>HSF: Creation of an exonic ESS site. Alteration of an exonic ESE site. Potential alteration of splicing. MaxEntScan: no result. SpliceAI: No impact (0.00)<br>Minigene: "No impact on splicing" | PM3_VS, PS3_sup, PM2, PP3, PP1_mod<br>**Pathogenic** | No change |
| CAPN3 NM_000070.2:c.2306G>A:p.(Arg769Gln) (hg19)chr15:42703124 Canonical Allele Identifier: CA127306<br>This variant has been previously identified in a homozygous and compound heterozygous state in multiple (>20) LGMDR1 patients, segregating with disease in several families (PMID: 7720071, 7762565, 9246005, 12461690, 14645990, 19015733, 16650086, 23553538, 30564623), (**PM3_VS, PP1_strong**). This variant is present at 1.5% frequency in Amish population (14/900 gnomADv3), due to the previously described founder effect (PMID: 9246005). Aside from this population, it is present at the MAX population frequency of 0.009% (4/42030 of African population, gnomADv3) (**PM2**). This variant is predicted to affect protein function (CADD 35, REVEL 0.947) (**PP3**). Functional studies in cells showed that this variant increase CAPN3 auto-catalytic activity (PMID 9642272) (**PS3**).<br>HSF: Activation of an exonic cryptic acceptor site, with presence of one or more cryptic branch point(s). Creation of an exonic ESS site. Potential alteration of splicing. MaxEntScan: impact on splicing. SpliceAI: Abnormal splicing (DS_DL=0.49) (**PP3**)<br>Minigene: "Mild impact on splicing" (**PS3_sup**)<br>Presence of an abnormal transcript lacking exon 22 (does not cause frameshift). The normal transcript (including exon 22) is expressed at similar level compared to control. Another abnormal transcript missing the last 87 nucleotides of exon 13 is present for both control and mutated constructs. | PM3_VS, PS3, PP1_Strong, PM2, PP3<br>**Pathogenic** | No change |
| CAPN3 NM_000070.2:c.2390A>C:p.His797Pro (hg19)chr15:42703494 Canonical Allele Identifier: CA392002246<br>This variant has been previously identified in a homozygous state in one LGMDR1 patient, with absence of 94kD CAPN3 protein on Western Blot (PMID: 17994539) (**PS3_mod, PM3_sup**). This variant is absent from population databases (gnomADv2.1.1 and v3) (**PM2**). This variant is not predicted to affect protein function (CADD 23,4, REVEL 0,528). It changes a non-conserved amino acid, located in the EF-hand domain of CAPN3 (PF13202).<br>There are two ClinVar entries for a different variant at the same position, c.2390A>G:p.His797Arg, classified as VUS.<br>HSF: Creation of an exonic ESS site. Alteration of an exonic ESE site. Potential alteration of splicing. MaxEntScan: no impact on splicing.<br>SpiceAI: No impact (0.01)<br>Minigene: "No impact on splicing" | PS3_mod, PM3_sup, PM2<br>**VUS** | No change |

## ABBREVIATIONS

ACMG: American College of Medical Genetics
aa: aminoacid
bp: base pair
CADD: Combined Annotation Dependent Depletion
CAPN3: Calpain 3
ESE: Exonic Splicing Enhancer
ESS: Exonic Splicing Silencer
HSF: Human Splicing Finder
kb: kilobase
kDa: kilodalton
LGMD: Limb Girdle Muscle Dystrophy
LOVD: Leiden Open Variation Database
RNA: ribonucleic acid

